# A critical role of *sux* cistron-mediated sucrose uptake for virulence of the rice blight pathogen *Xanthomonas oryzae* pv. *oryzae*

**DOI:** 10.1101/2025.06.02.657373

**Authors:** Nora R. Zöllner, Juying Long, Congfeng Song, Jacob Sharkey, Michael M. Wudick, Eliza P.I. Loo, Mayuri Sadoine, Violetta Applegate, Astrid Höppner, Sander H.J. Smits, Bing Yang, Wolf B. Frommer

## Abstract

The virulence of *Xanthomonas oryzae pv. oryzae*, the causal agent of bacterial blight (BB) of rice, critically depends on the activation of SWEET sucrose uniporters of the host. To date, the role of SWEET-released sucrose for virulence remains unclear. We here identified the *sux* locus of *Xoo*, consisting of a LacI-type repressor (SuxR), an outer membrane TonB-like porin (SuxA), an inner membrane MFS H^+^-symporter (SuxC), and a cytosolic sucrose hydrolase (SuxB). Structural and functional analyses demonstrate that SuxB has exclusive sucrose hydrolase activity. Mutant analyses show that the transporter SuxC and the sucrose hydrolase are necessary for growth of bacteria on sucrose, while SuxA is not essential, likely due to the ability of other porins to transport sucrose across the outer membrane. Consistent with a role of SuxR as a sucrose repressor, transcriptome studies show sucrose-dependent regulation of the *suxA/suxB* genes. Besides a role of sucrose for reproduction, we found that sucrose promotes motility, EPS production, biofilm formation, and virulence. Notably, the SuxC sucrose H^+^-symporter and the sucrose hydrolase SuxB were required for full virulence of *Xoo* on indica and japonica rice varieties. Our findings indicate that pathogen-induced sucrose efflux via SWEETs provides sucrose to *Xoo*, that *Xoo* uses the *sux* gene cluster to acquire and utilize sucrose, and that sucrose promotes bacterial fitness and xylem colonization.

**Significance Statement:** Understanding disease mechanisms is critical for developing strategies to protect plants against infections. Bacterial blight, a major threat to global rice production, is caused by *Xanthomonas oryzae pv. oryzae* strains that hijack rice SWEET sucrose uniporters. *Xanthomonas oryzae pv. oryzae* deploys a set of molecular “keys”, TAL effectors, to trigger host sucrose release. Sugars could serve as nutrients for the bacteria or prime host defense. Here, we provide evidence that the pathogen’s ability to attack and grow depends on sucrose uptake. We explore how crucial virulence functions, such as motility, EPS production, and biofilm formation, depend on the activity of the sucrose utilization system, supporting a key role of sucrose for the nutrition of the bacteria in the xylem.

## Introduction

Pathogens cause major crop losses, with massive economic and food security impacts. Bacterial blight (BB) is one of the major causes of rice yield losses, especially in Asia. Recent BB outbreaks in Tanzania and Madagascar and spread across East Africa led to substantial yield losses particulary affecting small-scale food producers (1). The causal agent of BB is the gram-negative bacterium *Xanthomonas oryzae pv. oryzae* (*Xoo*) (2). From the phyllosphere, *Xoo* invades the water-conducting xylem vessels by entering via wounds or hydathodes (3, 4). Within the xylem, *Xoo* moves in the xylem stream, then adheres to host cell walls, undergoing a transition from planktonic to sessile states from which new planktonic cells disperse and generate a progressive infection even against the xylem stream (3, 4). These transitions have to be timed precisely for effective infection. The regulatory networks driving infections of *Xoo* are highly complex, including light and quorum sensing, the secondary messengers c-di-GMP, and alternative sigma factors such as σ^54^ (5, 6). The regulatory network coordinates motility, biofilm formation, and virulence (7–9).

The xylem provides a niche with reduced microbial competition but presents major challenges, e.g., the rapid xylem flow (10, 11). Swimming against the stream is likely highly energy-intensive. The production of extracellular polysaccharides (EPS) and biofilm, required for withstanding the physical forces of the xylem flow, requires carbon skeletons and energy (12). Biofilm can protect against at least the loss of quorum-sensing molecules and metabolites, secreted in the newly forming colonies, fostering bacterial communication and reproduction (12–14). The high demand for carbon skeletons and energy poses major challenges since carbohydrate levels in the xylem sap are scarce, likely restricting bacterial virulence and growth (15–18). Freeze-dried guttation fluid, a proxy for xylem sap, failed to support *Xoo* growth, except when sucrose was added (19). *Xoo* metabolizes hexoses via the Entner-Doudoroff pathway, producing only one ATP compared to two ATP for the Embden-Meyerhof pathway, thus limiting ATP production from the low sugar available in the xylem sap (20, 21).

We hypothesize that *Xoo* utilizes sucrose as the major carbon and energy source during the infection. After adhering to the xylem walls, *Xoo* produces a type 3 secretion system (T3SS) to inject bacterial type III effectors (T3E) into the xylem parenchyma that induce *SWEET* sucrose uniporter genes (22, 23). As a consequence, SWEETs likely sequester sucrose, making it available to *Xoo* (24, 25). All pathogenic *Xoo* strains studied to date rely on activation of at least one of three sucrose uniporters (26–28). In contrast, glucose-transporting SWEETs do not support *Xoo* infection when artificially induced, and glucose uptake-deficient *Xoo* mutants remain able to colonize the xylem (28, 29). There are two hypotheses regarding the role of *SWEET* induction for pathogen resistance: host priming and pathogen feeding (30). For *Colletotrichum higginsianum* infections, *SWEET* induction may lead to an overaccumulation of sugars in the extracellular space, and high extracellular sugar levels have been implicated in defense priming via the salicylic acid pathway (31). In line with the priming hypothesis, sucrose supplementation led to increased resistance to blast (32). However, in the case of BB, the induction of *SWEETs* appears to be the basis of susceptibility, not increased resistance; resistance was increased when the bacteria could not induce a *SWEET*. The most parsimonious hypothesis for the role in BB is thus that carbohydrate levels in the xylem limit bacterial reproduction and virulence, and that sucrose released from the xylem parenchyma via SWEETs provides sucrose to the pathogen. The genomes of *Xanthomonads* showed an enrichment of transport systems that make use of TonB-like porins (TBDR)(33). Analysis of 76 TBDR mutants in *Xanthomonas campestris pv. campestris* (*Xcc*) identified the *TBDR XCC3358* gene as relevant for sucrose uptake. Deletion of the *TBDR XCC3358* gene caused delayed infection development (33). In *Xanthomonas axonopodis pv. glycines* (*Xag*) and *Xanthomonas axonopodis pv. manihotis* (*Xam*), mutations of respective sucrose utilization (*sux*) loci resulted in no or only moderate delay in symptom development, respectively (34, 35). To explore the role of sucrose for the virulence of *Xoo*, we tried to identify sucrose uptake systems. Here, we analyzed the *sux* cistron of *Xoo*, encoding a LacI-type repressor, an outer membrane TonB-like porin, an inner membrane MFS-family sucrose/H^+^ symporter, and a putative cytosolic amylosucrase. Transcriptomics were used to characterize the four genes in the *sux* operon. Structural, biochemical, and genetic analyses of the *sux* genes show that the *sux* locus is necessary for sucrose utilization in *Xoo* and plays critical roles for swimming, swarming, EPS production, and biofilm formation. Most importantly, the *sux* gene functions are critically required for virulence in japonica and indica varieties of rice. Sucrose-induced transcriptional regulation indicated that sucrose and its metabolic downstream products promote vital bacterial functions during xylem colonization, in particular after *SWEET* induction, as indicated by the presence of the *LacI-type repressor* gene in the cluster. We surmise that a better understanding of how host-derived sugars serve as nutrients for pathogens will help to develop innovative and long-term strategies against pathogen infection for BB, cassava, and cotton blight, but possibly also against other pathogens.

## Results

### Identification of a sucrose utilization gene cluster

To investigate whether and how *Xanthomonas oryzae pv. oryzae (Xoo)* can acquire sucrose, and to identify potential sucrose uptake and metabolic systems, we performed RNAseq on the Asian *Xoo* strain PXO99^A^. Cultures were grown *in vitro* in media supplemented with sucrose, glucose, or without additional carbon source, and samples were collected over an eight-hour time course. To assess transcriptomic changes and identify treatment-specific differences, we performed principal component analysis (PCA). PCA identified distinct temporal (PC1, 39.4 %) and treatment-specific (PC2, 20.1%) clustering. At 2 hours, the samples of sucrose, glucose, and without additional carbon sources clustered. At the four-hour timepoint, transcriptomes of the three media treatments diverged. Transcriptomes of samples grown without additional carbon sources became increasingly dissimilar from samples grown in sugar-supplemented media over time. (Fig. 1a).

**Figure 1.**
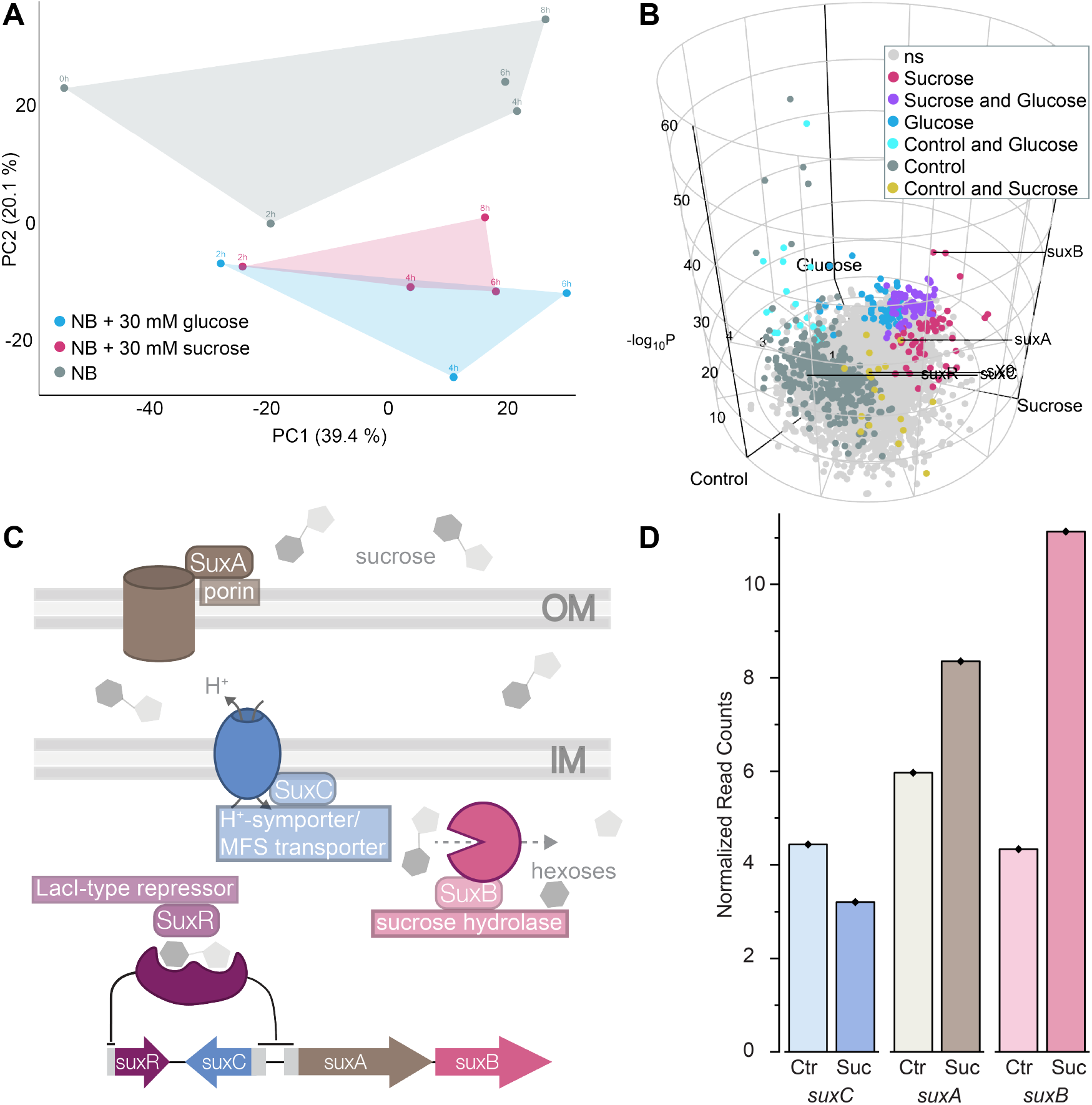
Identification of *sux* cluster in *Xoo* by RNAseq. PXO99^A^ cells were grown in NB media with and without supplementation of sugar and were sampled over eight hours for an *in vitro* RNAseq analysis. **A)** The first two principal components explaining the variance between *Xoo* grown in NB (grey), NB + sucrose (pink), and NB + glucose (blue). **B)** RNAseq data of *Xoo* grown for 4 hours in the three media types, where the abundance of each gene is assigned radial coordinates and color based on which treatment has more abundant RNA. The z-axis is the –log_10_ p-value with a significant p ≤ 0.01. **C)** Proposed model of the sux cluster-encoded sucrose uptake system in *Xoo*, including a schematic representation of the gene cluster. SuxA (brown): β-barrel outer membrane porin; SuxC (blue): major facilitator superfamily (MFS)-type sucrose/H^+^-symporter; SuxB (pink): cytosolic sucrose hydrolase; SuxR (purple): LacI-type transcriptional repressor. **D)** RNAseq read counts for *suxC* (blue), *suxA* (brown), and *suxB* (pink) after 6 hours of growth in media without additional sugar (Ctr) or supplemented with 30 mM sucrose (Suc). Values were normalized to the 0 h time point and the condition without additional sugar.

We identified 192 differentially expressed genes (DEGs) in sucrose-treated samples compared to no sucrose (p-value <0.05 and log2-fold change >2). Among the DEGs, 134 had decreased and 58 had increased steady-state transcript levels (Fig. 1b). The DEGs with increased transcripts were chaperones (*groE, clpB* and *htpG*), enzymes (*NADH-oxidoreductase*), membrane proteins such as *cytochrome B*, and an *outer membrane ß-barrel* transporter of the TonB family. Of particular interest regarding the acquisition of host-derived sucrose were two genes encoding an *amylosucrase* (PXO_02416, RS19450, hereafter named *suxB*) and an *outer membrane ß-barrel TonB-family transporter* (PXO_02415, RS19445, hereafter named *suxA*), both encoded in an operon encoding two proteins. To validate the sucrose-mediated induction of *suxAB* transcripts observed in the *in vitro* RNAseq data, bacterial growth assays were performed under various carbon supply conditions. The wildtype (PXO99^A^) exhibited minimal growth in the absence of an external carbon source, reflecting a base growth, whereas supplementation with sucrose or glucose elevated proliferation rates by 1.5-fold each (Fig. 2a). The coordinated induction in *amylosucrase* transcripts and the associated *outer membrane transporter* led us to hypothesize that *Xoo* evolved a distinct uptake mechanism for sucrose. SuxB showed 81 % protein identity to SuxB_*Xcc*_ in the *sux* genes from *Xanthomonas campestris pv. campestris* (33). Further analysis of the locus revealed a cistron composed of four genes including a *LacI-type repressor (suxR)* and an *inner membrane transporter* belonging to the major facilitator superfamily (MFS), likely functioning as a H^+^-symporter *(suxC)* (Fig. 1c; Supplementary Table 1). Notably, sucrose-induced DEGs from RNAseq included *cytochrome B* and *NADH-oxidoreductases*, both involved in the generation of the proton motive force (PMF), required for example for H^+^-symport and thus concentrative sucrose uptake (Supplementary Table 2).

**Figure 2.**
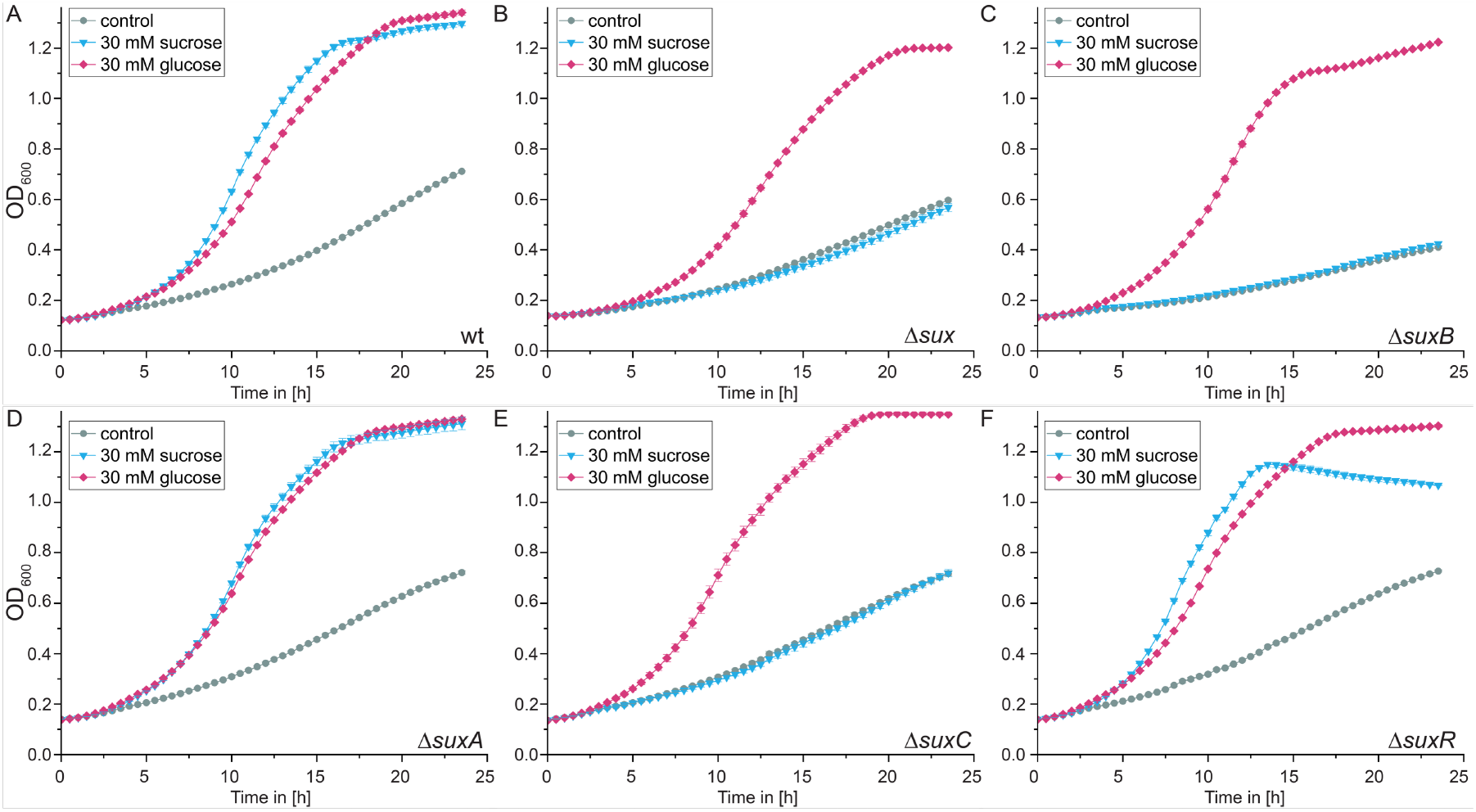
Effect of mutations in *sux* genes on bacterial growth. Growth curves of **A)** wildtype, **B)** *Δsux*, **C)** *ΔsuxR*, **D)** *ΔsuxC*, **E)** *ΔsuxA* and **F)** *ΔsuxB* in NB media (grey), supplemented with 30 mM sucrose (blue) or 30 mM glucose (pink). Mean of 3 technical replicates is shown with whiskers for ± SEM.

Analysis of *sux* gene clusters across 17 *Xanthomonads* revealed sequence identity >85 %. All 17 species, representing the African and Asian continents, with vascular and non-vascular lifestyles retained the *sux* cistron in the same organization (Supplementary Fig. 1). The cistron thus is a strong candidate for sucrose uptake and metabolism in *Xoo*. Temporal dynamics of transcriptional changes hypothesize a response of *Xoo* to sucrose, represented by the increase of *suxA* and *suxB* transcripts (Fig. 1d). The induction was sucrose-specific, as *sux* transcript levels remained unresponsive to glucose supplementation or showed a decrease (Supplementary Fig. 2). Changes in *suxR* transcripts were not observed. The conservation of the *sux* operon indicated a likely evolutionary importance, potentially unveiling a universal host-derived sucrose utilization mechanism in *Xanthomonads*.

### SuxB is a sucrose hydrolase

Using KEGG classification, SuxB was a member of family 13 of glycoside hydrolases and was classified as an amylosucrase (E.C. 2.4. 1.4), which either functions as a glycosyltransferase for synthesizing amylose-like polymers from sucrose or as a sucrose hydrolase. To differentiate between the activities, SuxB was crystallized in its apo state with a resolution of 2.4 Å (Fig. 3a). Independently produced crystals were soaked with sucrose, but likely due to hydrolytic activity, we identified a glucose molecule in the binding pocket of SuxB (resolution 2.95 Å, Fig. 3b). COOT was used to dock sucrose into the binding site based on the position of glucose (Supplementary Fig. 3). Structural analysis revealed a (β/α)_8_-barrel shape, which distinguishes SuxB from classical invertases with β-propellers linked to C-terminal β-sandwich structure (36, 37). SuxB lacks three arginine residues that are crucial for the glucosyltransferase activity of amylosucrases (37), indicating that SuxB is a sucrose hydrolase (GH13_4, E.C.3.2.1.48). The SuxB fold from *Xoo* is structurally similar to the sucrose hydrolase SUH_*Xag*_ from *Xanthomonas axonopodis pv. glycines* (37), with an average RMSD of around 0.6 over circa 535 Cα atoms. In both the apo and the glucose-complex structure, the asymmetric unit contains three SuxB molecules. When comparing the individual chains within the apo structure, all three SuxB molecules show identical arrangement with their corresponding chains with RMSD values ranging from 0.15 (over 590 Cα atoms, chains A) to 0.23 (over 582 Cα atoms, chains C). The structures differ only slightly for the conformations of the loops in the B and B’ domains. When using the Cα atoms of the residues His224 and Ser437 as reference points for describing the high degree of flexibility of the loops ‘shielding’ the active site (i.e. ∼ aa 221-225, loop between Bα3 and Bβ3; and ∼ aa 435-442, loop between B’β1 and B’β2) it becomes obvious: the distance between these two amino acids amount to 10.2 Å in chain A, to 7.7 Å in chain B and 7.5 Å in chain C in SuxB-apo (Fig. 3c). Beyond that, parts of these loops are described to not be visible in the electron density of the deposited structures of homologous proteins (37), which also indicates high flexibility in these parts.

**Figure 3.**
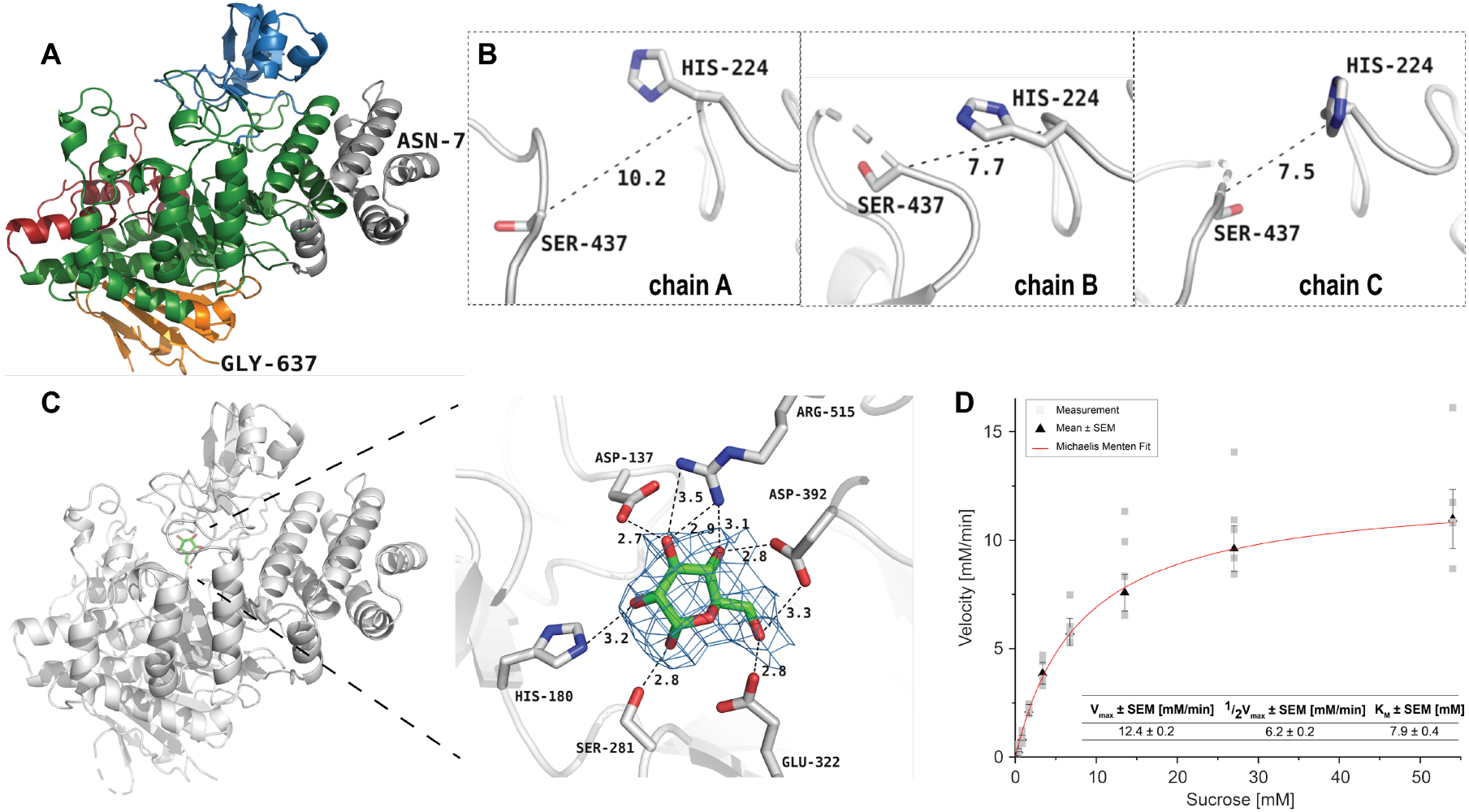
Characterization of *Xoo* sucrose hydrolase SuxB. **A)** Domains N, A, B, B’, and C are presented as cartoons accordingly to SUH (Kim et al., 2008) with domain N colored in grey, domain A in green, B in blue, B’ in red, and domain C in orange. For clarity, the N- and C-terminal residues Asn7 and Gly637 are labeled. **B)** The dashed box indicates the region of the flexible loops from domains B and B’, which are zoomed in. **C)** Overall structure of SuxB in complex with the hydrolase product glucose (left panel) and zoomed in on the active site (right panel). Highlighted are the active site residues that interact with the sugar molecule. The electron density around the bound glucose is contoured at 1 σ. In **A)** and **C)**, only chain A is presented. Additional data in supplementary figure 3. **D)** Michaelis-Menten curve of SuxB. Sucrose hydrolysis quantification via DNS assay on 0.1 µg/µL His-tag-purified enzyme in 20 mM MOPS, pH 7, at 28°C. (1/2) V_max_ = (half) maximum rate of reaction, K_M_ = Michaelis constant.

The accessibility of the active site was evaluated by analyzing the electrostatic surfaces, which show the widest opening in chain A and the closest access in chain B in the apo structures. Compared with the SUH_*Xag*_ structures (37), glucose in SuxB occupied the same position as the glucosyl moiety and the glucose molecule in SUH-E322Q-sucrose (pdb 3CZK) and SUH-glucose (pdb 3CZG) within the active site, with hydrogen bonds to the SuxB active site residues Asp137, His180, Ser281, Glu322, Asp392 and Arg515 (Fig. 3b). In contrast to the apo structure, all three chains in the SuxB-glucose complex appear to be more ‘rigid’ or stabilized by the interaction with the glucose, since the loops of the above-described B and B’ domains adopt a more similar conformation. AI prediction models support the hypothesis that SuxB recognizes sucrose and estimate the K_M_ to 5.5 mM and K_cat_ to 73.4 s^-1^ (38). To directly test for sucrose hydrolase activity, SuxB was expressed in *E. coli*. Colorimetric DNS (dinitrosalicylic acid) assays detected reduced sugars when sucrose was added (Fig. 3d). Further analysis revealed Michaelis-Menten kinetics with a K_M_ of 7.9 mM, similar to the predicted K_M_ (Fig. 3d). The identification of SuxB as a sucrose hydrolase strongly supports the hypothesis that the *sux* gene cluster is involved in sucrose utilization by *Xoo*.

### De-repression of sux genes by sucrose

The transcriptional regulator SuxR exhibits two conserved functional domains from the LacI family of bacterial regulatory proteins, including an N-terminal helix-turn-helix DNA binding domain and a C-terminal ligand binding domain (39). DNA binding motifs of SuxR were predicted using RegPrecise 3.0 and identified potential target regions in *PsuxC* and *PsuxAB* (Supplementary Fig. 4) (40). Consequently, we hypothesized that the ligand binding domain undergoes a conformational rearrangement upon sugar binding, which allosterically alters the affinity of the regulator to the DNA binding site. Motif analysis indicated that in the absence of sucrose, SuxR acts as a negative regulator (repressor) of the *sux* gene cluster. Regulation of SuxR might be mediated via the sigma factor σ^54^ (RpoN2). Promoter regions of *suxR* and *suxC* showed binding motifs for σ^54^, although imperfect for *suxR* (5, 41, 42) (Supplementary Fig. 4). In *Xoo*, motility, biofilm formation, EPS production, and virulence are controlled by the sigma factor σ^54^ (RpoN2). Based on the presence of the *LacI-type repressor suxR* in the cluster, and the ‘induction’ or derepression of *sux* genes by sucrose (Fig. 1d), we surmise that *in vivo*, the transcription activator-like effector (TALe)-induced SWEET activity will trigger release of sucrose that leads to derepression of the *sux* genes.

### sux gene functions are necessary for sucrose uptake

To evaluate whether the *sux* genes are required for sucrose utilization, deletion mutants were generated, either removing the whole *sux* gene cluster or deleting individual ORF encoding sequences. Deletion of the entire *sux* cluster *(Δsux*) abolished sucrose-dependent growth to base levels, while growth rates with or without glucose supplementation remained unaffected (Fig. 2b). Potential redundancies within the genome of the functions were tested using mutants for the individual genes in the cluster (Supplementary Fig. 4). The sucrose hydrolase SuxB was essential for sucrose utilization as indicated by a reduction in growth kinetics to base levels (Fig. 2c). The overall ability of the *ΔsuxB* mutant was unaffected, as indicated by having comparable growth rates as wildtype controls on media supplemented with glucose (Fig. 2c). SuxB is located downstream of SuxA, which is likely the porin responsible for the first step in sucrose uptake across the outer membrane (Fig. 1c). In contrast to the effect of *ΔsuxB* on sucrose utilization, *ΔsuxA* mutants did not show differences in growth kinetics when grown with sucrose or glucose (Fig. 2d). The well-established redundancy and overlapping substrate specificities of outer membrane porins or compensation might be possible explanations for the maintenance of sucrose transport across the outer membrane in the deletion mutant (33, 43, 44). Similar to *ΔsuxB*, the MFS transporter mutant *ΔsuxC* was unable to grow on media supplemented with sucrose but was characterized by similar growth rates as the wildtype in the presence of glucose (Fig. 2e). We did not expect that *ΔsuxR* would impair sucrose utilization. Consistent with this hypothesis, *ΔsuxR* was apparently unaffected and grew similar as wildtype (Fig. 2f). In summary, the results of the growth analysis of individual cluster components supported our hypothesis that sucrose uptake is facilitated by the *sux* cluster. Key players for sucrose uptake are *suxC* and *suxB*. Notably, glucose can act as an alternative carbon source to sucrose at least *in vitro*.

### Sucrose is necessary for EPS production and biofilm formation

Sucrose can be acquired via the *sux* gene cluster and serves as a carbon and energy source for reproduction. Carbon skeletons and energy are also required for biofilm production and to energize motility. Colony morphology can reveal deficiencies in EPS production and motility features. Phenotypic comparison with the wildtype revealed smaller colony diameters for *Δsux, ΔsuxC*, and *ΔsuxB* on media containing sucrose (Supplementary Fig. 5). Colony size on media without sucrose showed no difference from wildtype. Notably, *Δsux, ΔsuxC*, and *ΔsuxB* mutants appeared drier and flatter compared to wildtype on sucrose-supplemented media (Supplementary Fig. 5). *ΔsuxA*, lacking the outer membrane porin, did not show obvious deficiencies, as observed in the growth assays above (Fig. 3d). Supplementation with glucose rescued the morphology defects to wildtype levels (Supplementary Fig. 5). We hypothesize that *sux*-mediated uptake and metabolization by SuxB provides glucose as a precursor for EPS biosynthesis and biofilm formation. ‘Hanging drop’ assays on inverted plates containing sucrose showed that wildtype colonies slowly followed gravity in the form of a hanging drop, which eventually disintegrated, while *Δsux* colonies stayed relatively flat (Fig. 4a). Quantification of EPS by crystal violet staining showed a significant reduction in crystal violet staining between *Δsux* and wildtype cultures (Fig. 4b,c). The qualitative and quantitative data support the hypothesis that sucrose utilization in *Xoo* via the *sux* cluster is necessary for EPS production and biofilm formation, key factors required for virulence.

**Figure 4.**
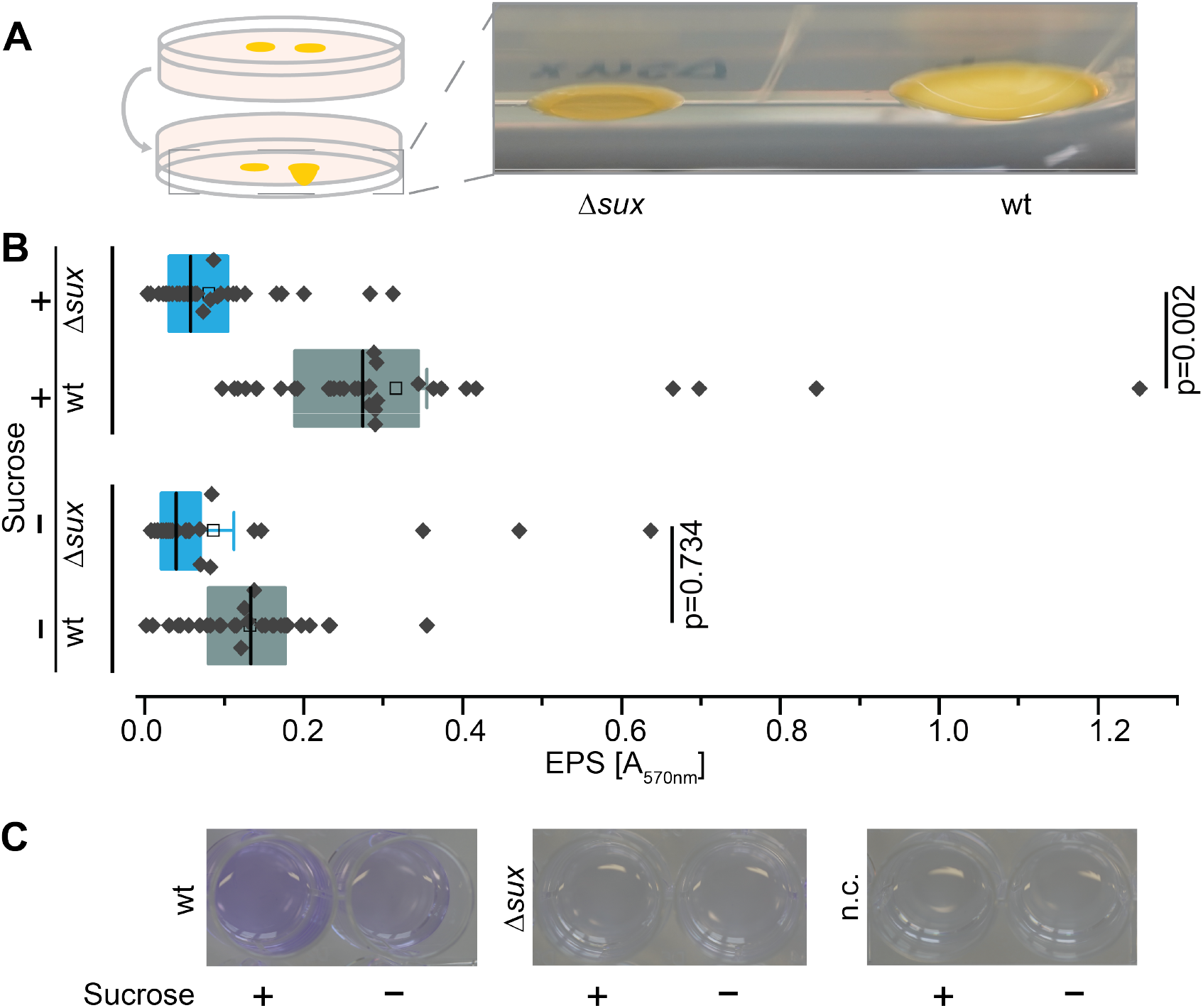
Effect of *sux* genes on EPS production and biofilm formation. **A)** Qualitative analysis of biofilm formation by inverted plates following the hanging droplet assay. Supplementary Figure 5 shows effects on glucose-supplemented media. **B)** Quantification of crystal violet from wildtype and *Δsux* cells in NB media ± 30 mM sucrose supply. Boxes range from 25th to 75th percentiles, with median values shown as center lines and mean values as empty boxes with whiskers ± SEM. Significance between two groups was calculated using unpaired two-tailed Student‘s t-test with 95 % confidence. **C)** Crystal violet staining of wildtype, *Δsux*, and negative control (no cells) in NB media ± 30 mM sucrose supply.

### sux gene function is necessary for swimming and swarming

We hypothesize that sucrose may also provide the energy required for motility during the progressive infection of the xylem against the stream direction. The energy demand for motility is primarily provided by PMF (45). *Xoo* transitions from planktonic to sessile lifestyle, marked by a shift from swimming to swarming. Swimming and swarming of *Δsux* was significantly reduced relative to wildtype (Fig. 5a, b). Together, our findings indicated that *sux*-mediated sucrose uptake drives motility in *Xoo*, likely by increased PMF generation.

**Figure 5.**
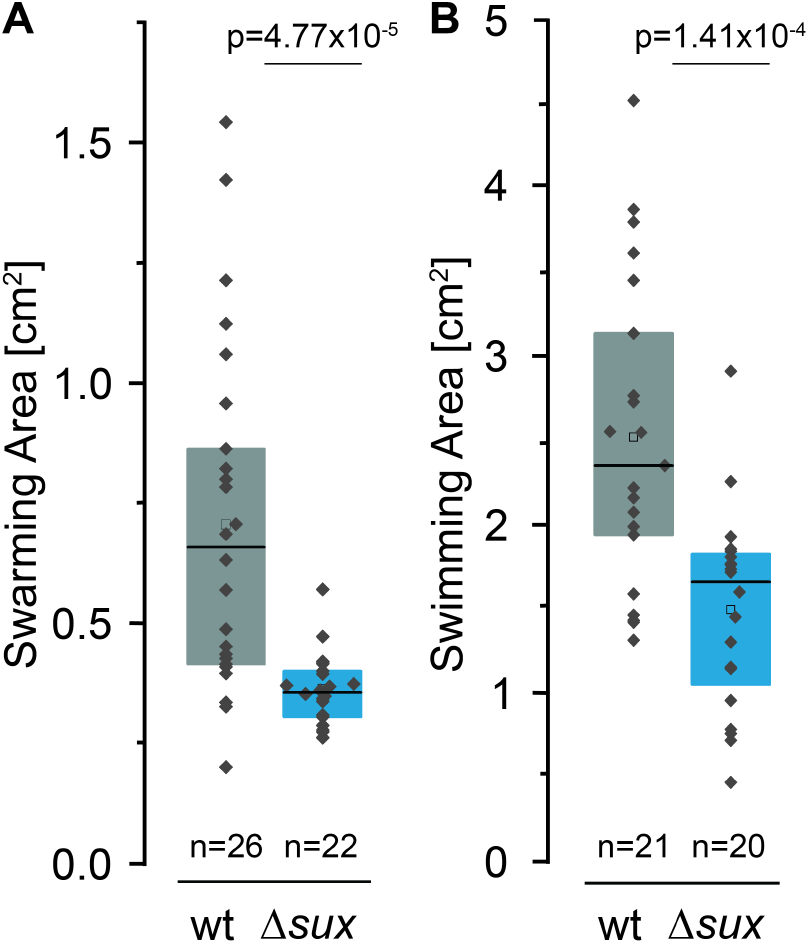
Role of *sux* genes in *Xoo* motility. Quantification of **A)** swarming area on 0.6 % agar plates and **B)** swimming area on 0.3 % agar plates of *Δsux* mutants (blue) compared to wildtype (grey). Boxes range from 25th to 75th percentiles, with median values shown as center lines and mean values as empty boxes with whiskers ± 1 SD. Significance between two groups was calculated using unpaired two-tailed Student‘s t-test with 95 % confidence.

### sux proteins are required for full pathogenicity

While the *in vitro* studies support a role of the *sux* gene cluster for sucrose uptake and virulence factor activity, a key question was whether the *sux* genes are also critical for successful infection of rice. To test for roles in virulence, three diverse rice cultivars, *Oryza sativa* ssp. *japonica cv. Nipponbare*, cv. *Kitaake*, and *Oryza sativa* ssp. *indica cv. IR24*, were infected with the *sux* mutants using clipping assays (46). Notably, virulence was significantly decreased in all three rice varieties (Fig. 6 a-d). Analyses of the contribution of the individual *sux* genes showed that while *ΔsuxA* was only slightly impaired, possibly due to TBDR redundancy, Δ*suxC* and Δ*suxB* mutants exhibited significantly reduced virulence (Fig. 6 a-d). Thus, the *sux* genes of *Xoo* play more critical roles for virulence than in *Xanthomonas axonopodis pv. glycines*, for which *suxB* mutation showed only moderate effects on virulence in soybean, or *Xanthomonas campestris pv. campestris*, where *sux* mutations only lead to a delay in symptom development on Arabidopsis (33, 34). Notably, the *sux* genes of *Xanthomonas axonopodis pv. manihotis* were not required for virulence in cassava (47). In summary, the *sux* genes of *Xoo* play a pivotal role in full virulence, linking *Xoo*-mediated transcription of the host *SWEET* transporters to SuxR-mediated derepression of a suite of *sux* genes crucial for uptake and utilization of sucrose, required for carbon and energy supply, as well as motility and adhesion.

**Figure 6.**
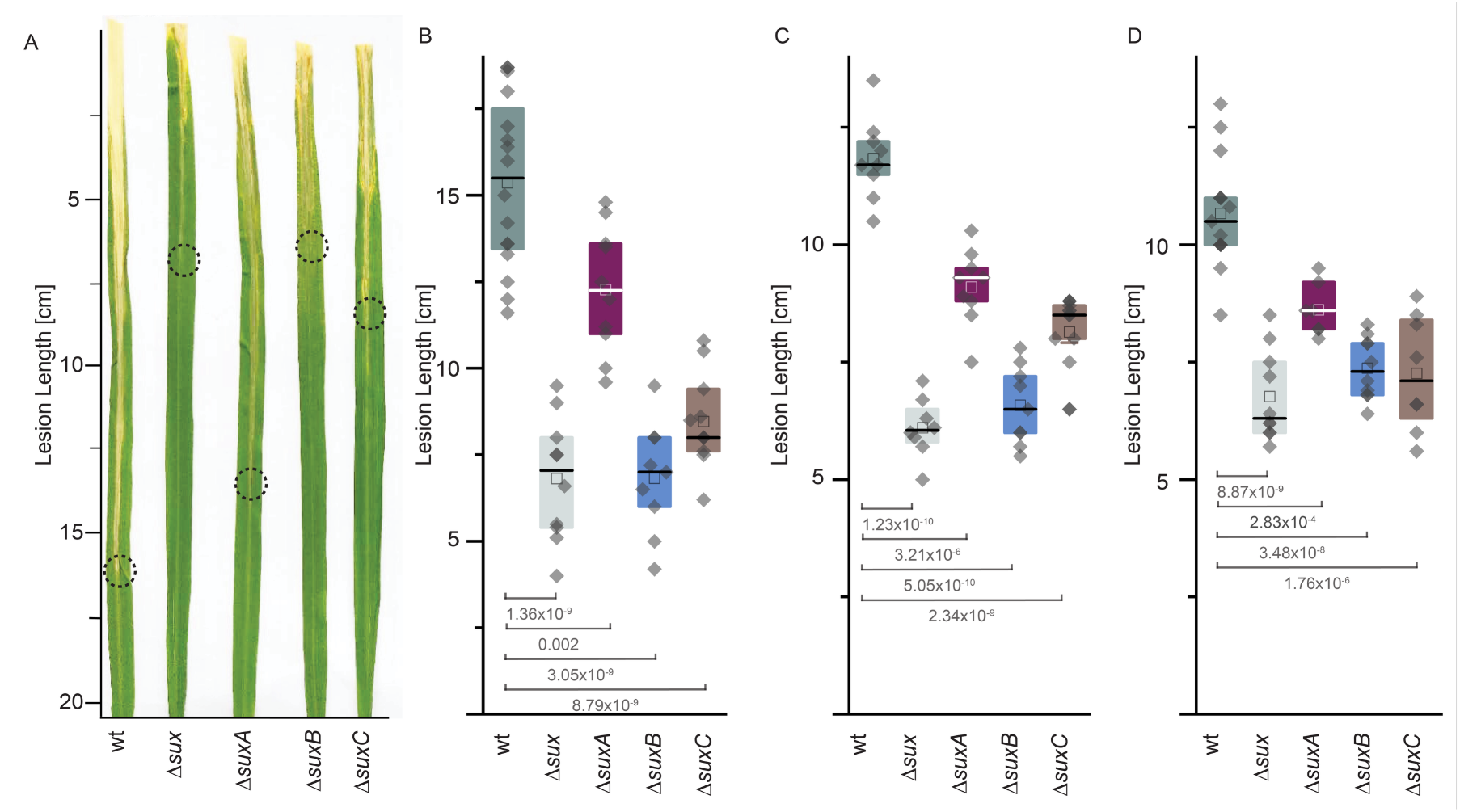
Impact of *sux* genes on *Xoo* virulence in rice bacterial blight. **A)** Representative leaf phenotypes in rice cultivar *Nipponbare* 12 days after infection with PXO99^A^ and *sux* mutants. **B-D)** Quantification of lesion length in rice cultivar **B)** *Nipponbare* (n=9-12), **C)** *Kitaake* (n=8-13), and **D)** *IR24* (n=7-15). Boxes range from 25th to 75th percentiles, with median values shown as center lines and mean values as empty boxes with whiskers ± SEM. Significance between two groups was calculated using unpaired two-tailed Student‘s t-test with 95 % confidence.

## Discussion

The dependency of *Xoo* virulence on TALe-mediated induction of a subfamily of sucrose-transporting *SWEETs* by *Xoo*, together with the uniport mechanism of SWEETs, and the low abundance of sucrose in the xylem of uninfected rice may intimate that the ability of *Xoo* to colonize the xylem depends critically on sucrose supply. Here, we identified the *sux* cistron as an essential cluster of genes required for sucrose uptake and utilization in *Xoo*. Importantly, the *sux* gene cluster is required for virulence, and *sux* mutants can cause only low levels of disease, indicating that sucrose is a key limiting factor for successful colonization.

*Xoo* likely uses carbon sources other than sucrose while living outside a plant, as well as in early stages of infection of the plant, when sucrose is not present and the *sux* genes are repressed. We hypothesize that upon release of sucrose due to induction of *SWEET*s, *sux* genes are derepressed, enabling the bacteria to acquire and metabolize sucrose. The key components of the cistron include a LacI-type repressor SuxR, and sucrose induction of the *sux* genes supports a sucrose-dependent derepression of the genes. As a γ-proteobacterium, *Xoo* has to take up nutrients across the outer and inner membranes. The porin-like SuxA likely mediates uptake across the outer membrane. However, Δ*suxA* mutants could still grow on media with sucrose, and virulence was only partially inhibited. Other porins either compensate or can mediate sufficient uptake of sucrose across the outer membrane (33). In contrast, the MFS-family transporter SuxC, which likely functions as a secondary active H^+^-symporter on the inner membrane, was essential for growth on sucrose-containing media and showed a similar reduction in virulence as mutants carrying deletions of the complete gene cluster. At the same time, sucrose triggers an increase in transcripts for proteins involved in generating the PMF, namely Cytochrome B and NADH-oxidoreductases, required for H^+^-symport and thus concentrative sucrose uptake. The fourth gene in the *sux* gene cluster, *suxB*, encodes a protein annotated as amylosucrase, so an enzyme that can hydrolytically cleave sucrose and synthesize amylose-like polymers from sucrose. Analysis of crystal structures supports the hypothesis for hydrolysis activity based on the apo-, glucose-bound, as well as modelled sucrose-bound structures. The active site of SuxB is similar to the SUH protein from *Xag*, and the binding site for the fructosyl and glucosyl moieties corresponds to the +1 and −1 subsites, respectively (48). Three arginine residues were crucial for glucosyltransferase activity of amylosucrases of *Neisseria* (34). Mutant forms of the sucrose hydrolase from *Xag* failed to restore glucosyltransferase activity (37). The arginine residues are not conserved in SuxB from *Xoo*, strongly supporting a sole function as sucrose hydrolase. Kinetic analysis shows that SuxB has a K_M_ of 7.9. If we assume that the K_M_ is indicative of the sucrose levels, and given the potential dilution by the xylem flow, the value appears relatively high, at least in early phases where single *Xoo* cells inject TALes to trigger sucrose efflux. H^+^-symport by SuxC may enable concentrative sucrose uptake at very low supply levels. The phenotypical analysis of the mutants demonstrates that sucrose utilization is critical for growth as well as motility, EPS production, and biofilm formation. In summary, the data support a role of the *sux* gene cluster as the key uptake and utilization system for *Xoo*. Notably, the mutation of the *sux* gene cluster leads to a significant reduction in virulence. Natural infections are initiated by a single or a few bacterial cells that multiply and cause disease. In contrast, virulence assays using the clipping method rely on high bacterial titers for inoculation, which may not reflect natural infection dynamics (49). The residual virulence observed in this study may therefore not be relevant under natural conditions. Nevertheless, the PXO99^A^ strain lacking the TALe *PthXo1*, which induces *SWEET11a*, is avirulent (50). While *sux* mutants exhibit substantially reduced virulence, their ability to sustain a low level of pathogenicity indicates that alternative carbon sources might compensate. SWEET-derived sucrose may also be hydrolyzed in part by secreted host cell wall invertases, e.g., *OsCIN1, OsCIN4*, and *OsCIN7*, for which transcript levels increase during *Xoo* infection (51).

### Potential relevance of the role of nutrient supply beyond BB

γ-proteobacteria, a class of the proteobacteria to which *Xanthomonads* belong, includes many important pathogens, e.g. *Yersinia, Escherichia or Legionella*, that cause disease in animals and humans; or the genus *Xanthomonas*, comprising many pathogenic bacteria causing disease on a wide spectrum of plants, and *Pseudomonas*, which can infect animals and plants or be beneficial (52). Notably, many bacterial genera, including γ-proteobacteria, can cause disease in animals and plants, e.g., *Salmonella, Shigella, Enterobacter, Enterococcus, Pantoea, Burkholderia, Rhizobium*, or *Pseudomonas. Xanthomonads* are predominantly known as plant pathogens causing diverse diseases across a wide range of plants and including major crop diseases. There are indications that *Xanthomonas* has been found in human blood and nosocomial patients have been found. The identification of disease mechanisms and fundamental dependencies of *Xanthomonads* may have relevance beyond a single species, such as *Xoo* here, but possibly across multiple species and possibly other genera within the γ-proteobacteria. Notably, the TALe-mediated induction of *SWEETs*, found originally in *Xoo*, also appears to be important for cotton and cassava blight (47, 53). We surmise that pathogens infect hosts primarily to gain access to the host’s nutrient pool as an essential prerequisite for reproduction, as well as a source of energy for virulence-related functions. Genome editing has enabled the development of resistant varieties, and regulatory pipelines have been implemented to assess performance and transgene/foreign DNA elimination by segregation in accordance with national biosafety standards (54). Such tools lay the groundwork for future crop protection strategies. Combined with a comparative characterization of the metabolic functions that relate to virulence, may provide new ways to protect hosts from infections or cure diseases.

## Methods

### Bacterial strains, plasmids, and DNA constructs

One Shot TOP10 *Escherichia coli* cells (ThermoFisher, C404006) were cultivated at 37°C in LB (5 g/L yeast extract, 10 g/L tryptone, 10 g/L NaCl, pH 7). When required, kanamycin (50 µg/mL) was added for selection. *Xanthomonas oryzae pv. oryzae* strain PXO99^A^ was grown at 28°C in optimized nutrient broth (NB) (1 g/L yeast extract, 3 g/L beef extract, 5 g/L peptone, pH 7, 1.5% w/v agar) (55). When necessary, medium was supplemented with sugar (30 mM sucrose or 30 mM glucose) or antibiotics (25 µg/mL kanamycin or 50 µg/mL spectinomycin). Standard DNA techniques were used for *E. coli* and recombinant DNA manipulations as previously described (56).

### Construction of *sux* mutants in PXO99^**A**^

Homologous recombination was used to generate *Xoo* mutant lines. Genomic DNA (gDNA) of PXO99^A^ was extracted using the DNeasy Blood & Tissue kit (Qiagen, 69506). Flanking regions of roughly 200 bp up- and downstream of the gene of interest were amplified from gDNA using restriction digest cloning. The flanking sites were cloned in-frame around a spectinomycin cassette into the suicide vector pK18sB in *E*.*coli* cells. pK18sB was a gift from Gredd Beckham and Christopher Johnson (57). Electrocompetent PXO99^A^ was transformed with 500 ng of plasmid DNA via electroporation for 5 sec at 2.5 kV and 200 Ω in a 0.1 cm MicroPulser cuvette (Bio Rad, 1652089). After recovery in Recovery Medium For Expression (Sigma, CMR001-8×12ML), sucrose-tolerant, spectinomycin-resistant, and kanamycin-sensitive colonies were selected in multiple rounds of re-plating.

### Bacterial growth studies of PXO99^A^ and mutant strains

Bacterial growth studies of PXO99^A^, *Δsux, ΔsuxR, ΔsuxC, ΔsuxA*, and *ΔsuxB* were initiated at a starting OD_600_ of 0.1-0.2 using a mid-logarithmic pre-inoculum. Cultures were grown in 12-well, flat-bottom, transparent plates in a plate reader (Tecan Spark) at 28°C and 200 rpm of constant shaking. *Xoo* was grown in NB, NB + 30 mM sucrose, and NB + 30 mM glucose. OD_600_ was measured every 30 minutes over a course of 24 hours.

### Total RNA isolation and transcriptome analysis

100 mL media (NB, NB + 30 mM sucrose, and NB + 30 mM glucose) were inoculated in Erlenmeyer flasks with a mid-logarithmic pre-inoculum of PXO99^A^ to OD_600_ of 0.1-0.2. Cultures were incubated on an orbital shaker at 28 °C and 200 rpm. OD_600_ was measured every two hours over a period of 8 hours. Additionally, triplicates of 2 mL samples were taken and total RNA was extracted using the TRIzol™ Max Bacterial RNA Isolation kit (ThermoFisher, 16096020) followed by Direct-zol RNA Miniprep kit (Zymo, R2051). Complementary DNA (cDNA) library prep for Illumina sequencing was performed on pooled triplicates of total RNA by Novogene (UK, Cambridge). For qRT-PCR, total RNA was transcribed into cDNA using the Maxima H kit (ThermoFisher, M1682). 2 ng of cDNA were used as template. The 2^-ΔΔ^ct method was performed, and Δct values are presented in the analysis with *gyrA* as housekeeping gene (58).

### Differential Expression Analysis and Visualization

Raw gene counts were quantified by Novogene with FeatureCounts (82) and normalized to counts per million (CPM) using EdgeR. Principal component analysis (PCA) was completed with the prcomp() function of the stats R package. The final PCA plot was generated in R with ggplot2. Overlap of treatment groups is visualized by overlaying the PCA plot with convex hulls using the chull() function of the grDevices package (83). Differential expression was visualized using a modified version of the volcano3D package (84). Gene length-corrected trimmed mean of M-values was calculated for each sample using raw gene counts as input (85). P-values and fold changes were determined at each time point by comparing three media types (NB, NB + glucose, and NB + sucrose). P-values, fold changes, and z-scaled expression values were input into the polar_xy() and polar_p() functions of the volcano3D package, producing polar coordinate tables formatted for the volcano3D plot. The RNAseq data and R scripts used in these visualizations are deposited in the DataPlant ARC, published alongside this paper.

### Crystallization, data collection and structure refinement for SuxB-apo and SuxB in complex with glucose

Initial screening for crystallization conditions was performed at 12 °C in sitting drop plates using vapor diffusion method and commercially available screens. Crystals were optimized and appeared at 0.25 M potassium sodium tartrate and 32.5 – 37.5 (w/v) % PEG 3350 as precipitant. Before crystals were harvested for data collection, 1 µL ethylene glycol was added to the crystallization drops, and these were overlaid with mineral oil as a cryoprotection protocol before they were flash frozen in liquid nitrogen. A complete data set for the SuxB apo structure was collected at the ESRF (Grenoble, France), beam line ID23-2, at 100 K. Data were auto-processed with EDNA_proc and together with an AlphaFold3 model of SuxB used in molecular replacement for solving the structure (phaser_MR, ccp4 suite). The SuxB structure was built further and refined using iterative cycles of manual building in COOT and automated refinement processes using PHENIX Refine. Data collection and refinement statistics can be found in Supplementary table 3. Crystallization of SuxB in complex with glucose was performed as described above, but cryoprotection procedure was slightly different: Before crystals were harvested, 1 µL of a saturated sucrose solution was added to the crystallization drops before they were overlaid with mineral oil and flash frozen in liquid nitrogen. A complete data set for the SuxB complex structure was collected at the EMBL, beam line P13 (DESY, Hamburg, Germany) and auto-processed with autoPROC. Molecular replacement with phaser_MR was performed using the refined structure of apo SuxB. Model building and refinement were performed as described above, additionally, in two of the three chains of SuxB within the asymmetric unit, could be manually fitted to one fructose molecule in the additional density at the active site.

### Monitoring bacterial colony morphology

To assess phenotypic characteristics, 3 µL of cultures from deletion strains or wild-type PXO99^A^ were spotted onto solid NB agar with or without 30 mM sucrose. Photos were captured 3 days after incubation at 28 °C.

### Analysis of biofilm formation

Biofilm formation was evaluated using a ‘hanging droplet’ assay. After culturing *Xoo* on solid NB media with 30 mM sucrose, plates were inverted so that the agar was on top. Plates were left to stand for 30 minutes. The presence or absence of hanging biofilm droplets was observed and photographed.

### Motility assays

*Xoo* was grown in NB media in Erlenmeyer flasks to mid-logarithmic phase before dilution to a starting OD_600_ of 0.1. 2 μL of the dilution were dropped onto 0.6 % agar plates with 30 mM sucrose for swarming assays, and onto 0.3 % agar plates with 30 mM sucrose for swimming motility assays. Motility areas were measured 3 days after incubation at 28 °C.

### Analysis of EPS production

*Xoo* was grown in NB medium in Erlenmeyer flasks to mid-logarithmic phase before dilution to a starting OD_600_ 0.1-0.2 in NB media with 0 mM sucrose, 30 mM sucrose, or 30 mM glucose. Cells were grown for 24 hours at 28 °C on an orbital shaker at 200 rpm in 24 well-plates, medium without *Xoo* served as control for background subtraction. Plates were removed from the shaker and kept standing still for 24 hours before quantifying EPS using crystal violet (CV) staining. Medium was removed, and wells were washed with dist. water. 0.1 % CV (certified by the Biological Stain Commission, Sigma) was added to the wells and incubated for 10 minutes before rinsing with water. Finally, bound CV was solubilized in ethanol. Plates were photographed before CV quantification in a spectrophotometer (emission at 570 nm).

### Disease assays in rice leaves

*Oryza sativa ssp. japonica cv. Kitaake, cv. Nipponbare*, as well as *Oryza sativa ssp. indica cv. IR24* were grown in a plant growth chamber (12 h light, 28 °C; 12 h dark, 25 °C, 80 % humidity). Four-week-old rice plants were inoculated with bacterial suspensions in sterile distilled water at approximately OD_600_ 0.5 using leaf tip-clipping. Lesion lengths were quantified 12 DAI as described (59).

### Building the phylogenic tree of the *sux* gene cluster

Sequences were aligned using MAFFT and curated with BMGE to remove poorly aligned regions. A maximum likelihood phylogenetic tree was inferred with PhyML, using the GTR substitution model. The tree was rendered and displayed in Newick format with 10000 replicates for bootstrapping and visualization using http://www.ngphylogeny.fr (60).

## Statistics

Statistics were performed using the OriginLab Stats Adviser (OriginPro 2024b).

## Acknowledgments

We thank Susanne Paradies for excellent technical assistance. We would like to thank Boris Szurek for advice on *Xoo* transformation. We would like to thank Eduardo Patriarca (CNR-IGB, Naples, Italy) for the “hanging droplet assay”. This work was supported by grants from Deutsche Forschungsgemeinschaft (DFG, German Research Foundation) - Collaborative Research Center SFB1535, project ID 458090666/CRC1535/1; Deutsche Forschungsgemeinschaft (DFG, German Research Foundation) under Germany’s Excellence Strategy – EXC-2048/1 – project ID 390686111, Bill and Melinda Gates Foundation to HHU (WBF), with a subcontract to MU (BY) (INV-008733, INV-063189); and the Alexander von Humboldt Professorship. The Center for Structural Studies is funded by the Deutsche Forschungsgemeinschaft (DFG Grant number 417919780, INST 208/740-1 FUGG, INST 208/868-1 FUGG, and INST 208/761-1 FUGG) and Deutsche Forschungsgemeinschaft (DFG, German Research Foundation) – SFB1535 - Project ID 458090666 Project Z01 (SHJS & WBF).

