## Supplementary for "A critical role of *sux* cistron-mediated sucrose uptake for virulence of the rice blight pathogen *Xanthomonas oryzae* pv. *oryzae*"

**Supplementary table 1. Genes of the *sux* locus in *Xoo* with KEGG-predicted function**

| Locus ID PXO99[A] | | UniProt ID | Gene | Putative function |
|---|---|---|---|---|
| PXO_02412 | PXO_RS19435 | A0A0K0GP99 | *suxR* | sucrose repressor (LacI-type transcriptional regulator with HTH* domain) |
| PXO_02413 | PXO_RS19440 | A0A0K0GPI3 | *suxC* | sucrose/$H^+$-symporter (inner membrane MFS**-type sugar transporter) |
| PXO_02415 | PXO_RS19445 | A0A0K0GNQ5 | *suxA* | β-barrel outer membrane porin |
| PXO_02416 | PXO_RS19450 | A0A0K0GP78 | s*uxB* | amylosucrase of GH13 family*** (α-amylases), sucrose hydrolase |

* Helix-Turn-Helix
** Major Facilitator Superfamily
*** Glycoside Hydrolase Family 13

**Supplementary table 2. RNAseq PMF candidates**

| gene_id | gene_id (RS) | log2Fold Change | P value | padj | gene_description |
|---|---|---|---|---|---|
| PXO_03768 | PXO_RS02020 | 2.53 | 0.041 | 0.995 | NADH oxidase |
| PXO_01688 | PXO_RS14615 | 2.92 | 0.025 | 0.995 | Cytochrome b |

**Supplementary table 3. Data collection and refinement statistics.**

| | **SuxB_apo** | **SuxB_glucose** |
|---|---|---|
| Data collection | ESRF, ID23-2<br>25.01.2025 | DESY, EMBL; P13<br>22.03.2025 |
| Wavelength [Å] | 0.873130 | 1.059700 |
| Resolution range [Å] | 46.98 - 2.40 (2.49 - 2.40) | 49.75 - 2.95 (3.06 - 2.95) |
| Space group | I 2 2 2 | I 2 2 2 |
| Unit cell [Å, °] | 98.754 175.959 258.335<br>90 90 90 | 100.535 172.726 259.777<br>90 90 90 |
| Total reflections | 175616 (17385) | 632095 (63699) |
| Unique reflections | 88004 (8702) | 47939 (4747) |
| Multiplicity | 2.0 (2.0) | 13.2 (13.4) |
| Completeness (%) | 1.00 (1.00) | 1.00 (1.00) |
| Mean I/sigma (I) | 7.95 (1.75) | 11.07 (1.65) |
| Wilson B-factor [Å$^2$] | 33.76 | 91.52 |
| R-merge | 0.08 (0.41) | 0.13 (1.39) |
| CC1/2 | 0.99 (0.72) | 1.00 (0.88) |
| CC* | 1.00 (0.92) | 1.00 (0.97) |
| Reflections used in refinement | 87997 (8700) | 47909 (4744) |
| Reflections used for R-free | 4418 (443) | 2465 (272) |
| R-work | 0.17 (0.23) | 0.24 (0.47) |
| R-free | 0.23 (0.31) | 0.30 (0.51) |
| CC (work) | 0.97 (0.90) | 0.93 (0.12) |
| CC (free) | 0.94 (0.76) | 0.92 (0.10) |
| Number of non-hydrogen atoms | 15377 | 14527 |
| macromolecules | 14483 | 14484 |
| Ligands | 154 | 38 |
| Protein residues | 1861 | 1864 |
| RMS (bonds) | 0.01 | 0.01 |
| RMS (angles) | 1.26 | 1.26 |
| Ramachandran favored (%) | 97.40 | 94.00 |
| Ramachandran allowed (%) | 2.49 | 5.35 |
| Ramachandran outliers (%) | 0.11 | 0.65 |
| Rotamer outliers (%) | 0.07 | 12.00 |
| Clashscore | 7.16 | 7.98 |
| Average B-factor | 41.05 | 118.92 |
| macromolecules | 41.08 | 118.97 |
| ligands | 50.98 | 105.31 |
| solvent | 38.50 | 83.12 |
| PDB code | 9R8I | 9R9L |

Refinement parameters are highlighted. Statistics for the overall data quality and for the highest-resolution shell these are shown in parentheses. Statistics for the highest-resolution shell are shown in parentheses.

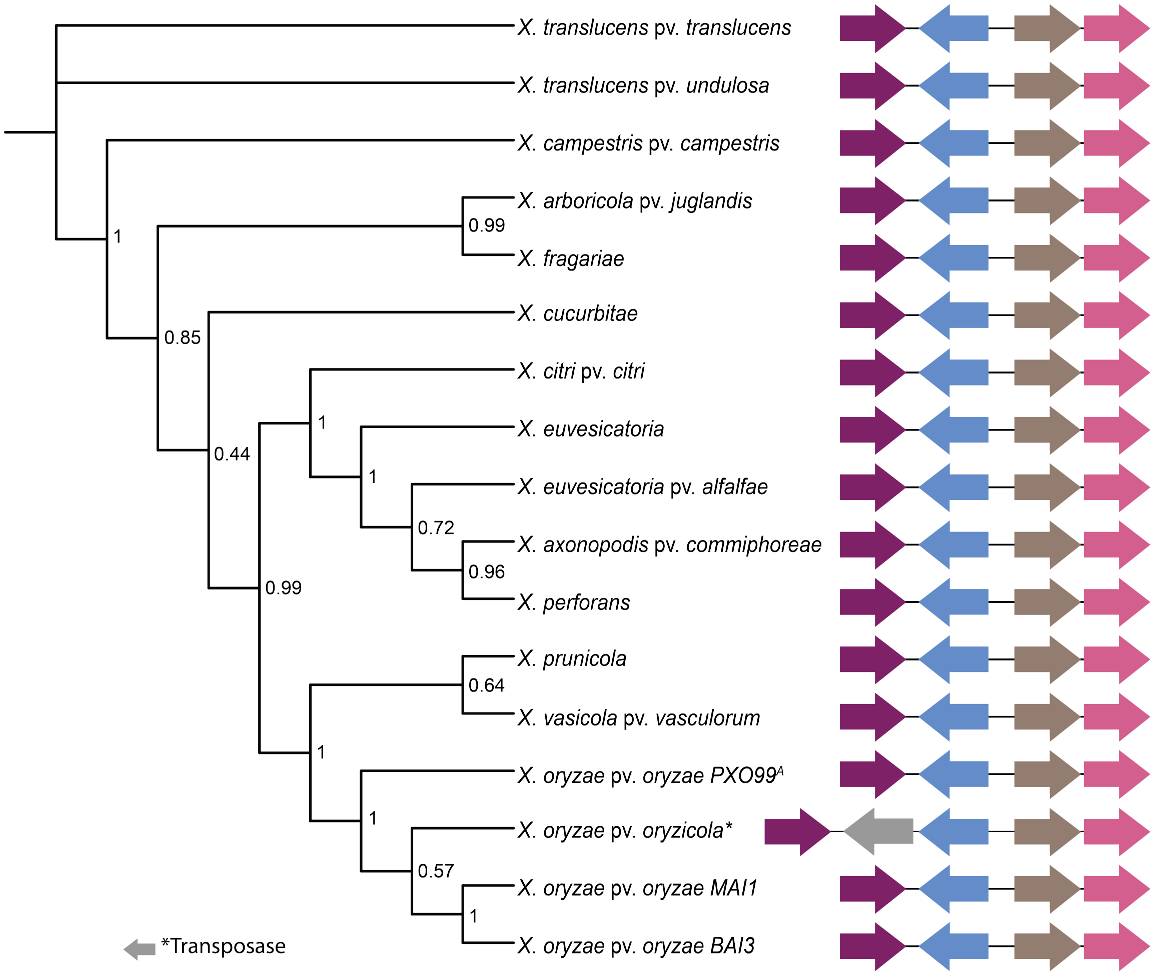


**Supplementary figure 1. Phylogenetic tree of *sux* cluster in *Xanthomonas* species.** Homologous genes based on *Xoo* PXO99$^{A}$'s *sux* cluster. The analysis spans 17 *Xanthomonas* species from strains of the African and Asian continents with vascular and non-vascular lifestyles. Depiction of conserved cluster structure with *suxR* (purple), *suxC* (blue), *suxA* (brown), and suxB (pink), as well as an identified *transposase* gene (grey) in *Xoc*. Phylogenetic confidence given by bootstrap values at nodes.

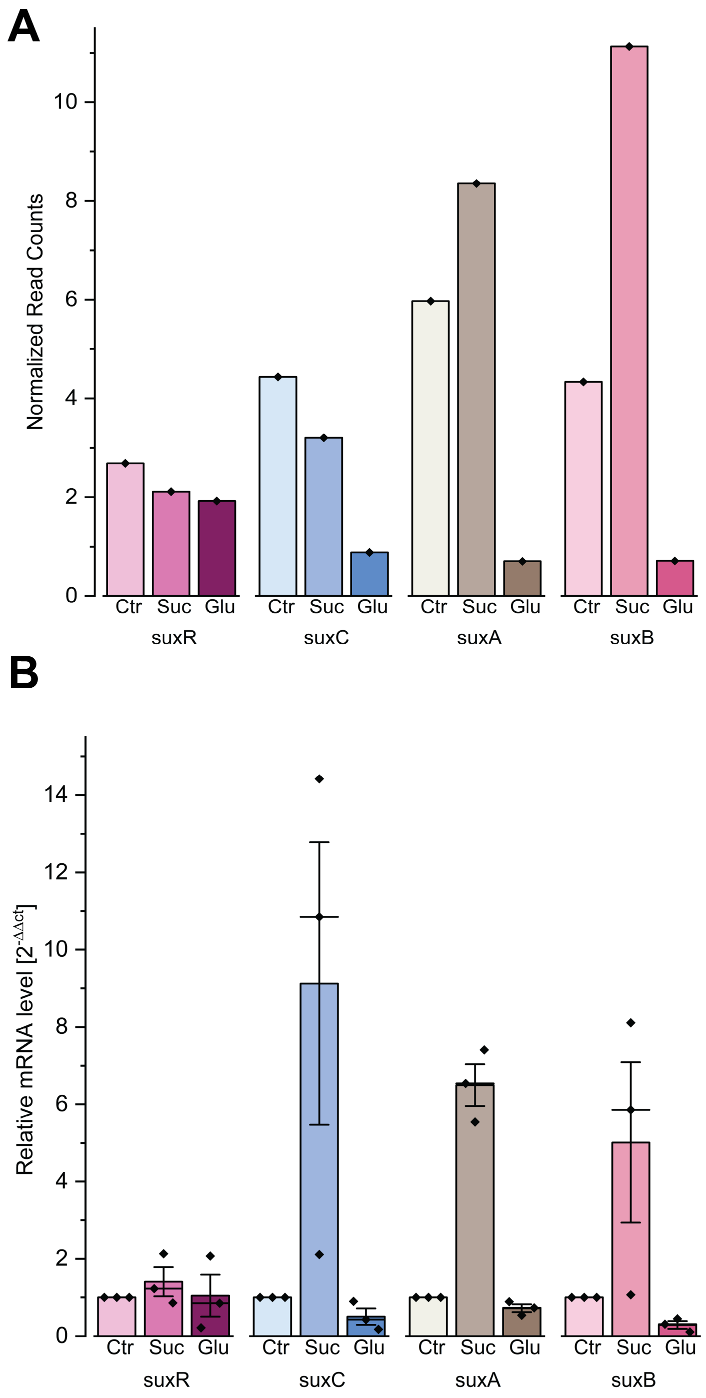


**Supplementary figure 2. Transcriptomic analysis of the *sux* cluster.**
Transcriptomic analysis of all *sux* gene mRNA levels. Cells were grown in NB media without added sugar (Ctr) or with 30 mM sucrose (Suc) or glucose (Glu). **A)** Read counts from the 6h time point of the RNAseq data set were normalized to the untreated condition and 0h time point. The trends were compared to **B)** qRT-PCR results of the 6h time point. Normalization based on $2^{-\Delta\Delta ct}$ method by (80). Display of three data points from two independent experiments. Bar height represents mean, horizontal line represents median, error bars display ± SEM.

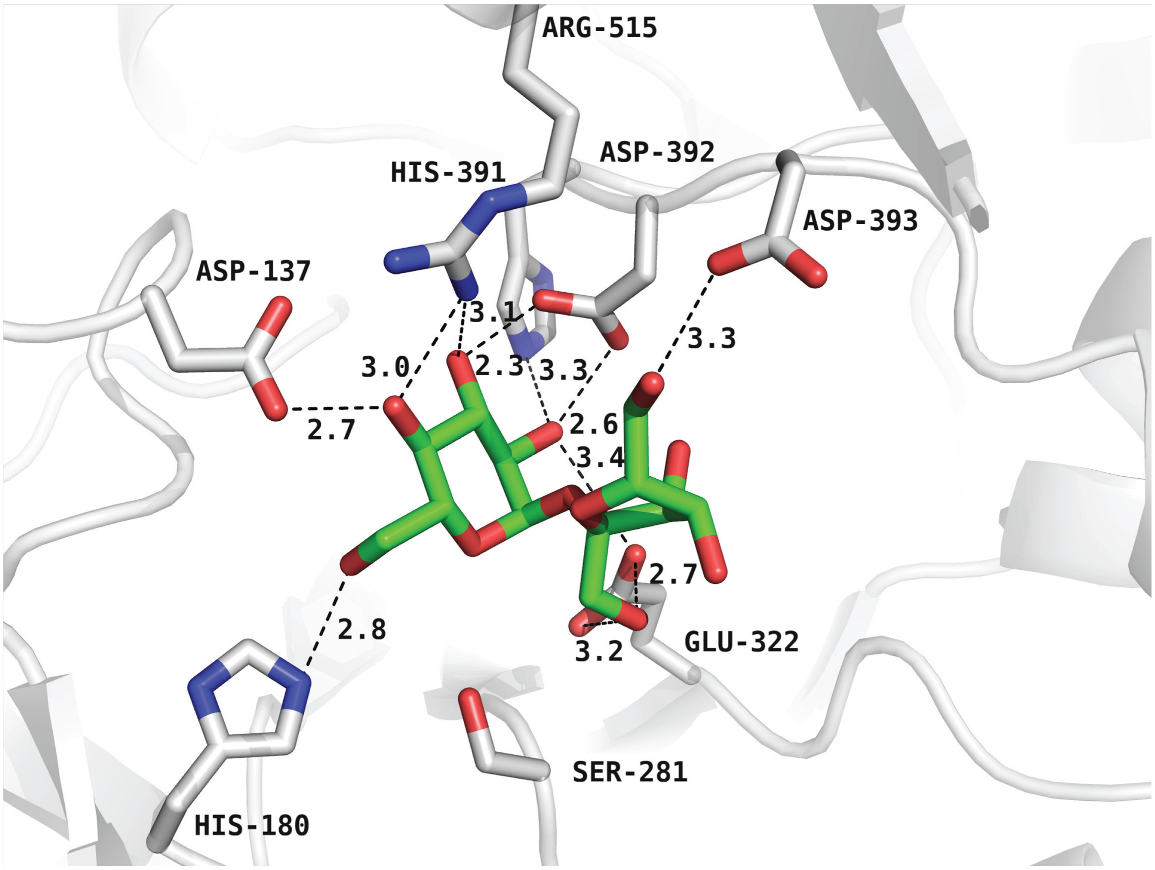


**Supplementary figure 3. Sucrose binding site of SuxB (glucose-bound structure PDB: 9R9L).** Zoom in to the active site of SuxB with a sucrose molecule fitted in manually using COOT and the position of the glucose molecule as anker point. The glucose moiety is bound via interactions with the side chains of Asp137, His180, His391, Asp392 and Arg515, whereas the fructose moiety interacts with Glu322 and Asp393.

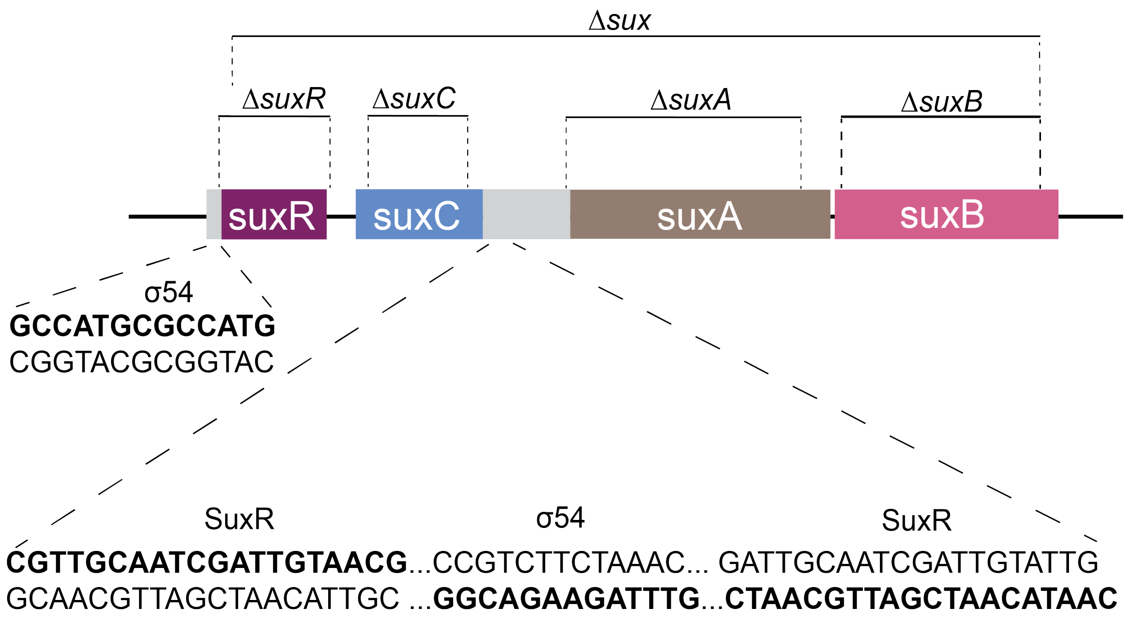


**Supplementary figure 4. Mutants used in this study and binding motifs.**
*Sux* cluster structure with marked regions (dotted lines), which were replaced by a spectinomycin cassette during mutant generation. Promoter regions are colored in grey, *suxR* in purple, *suxC* in blue, *suxA* in brown, and *suxB* in pink. Putative binding sites for σ54 transcription factor and SuxR are highlighted and bolded for strand specificity.

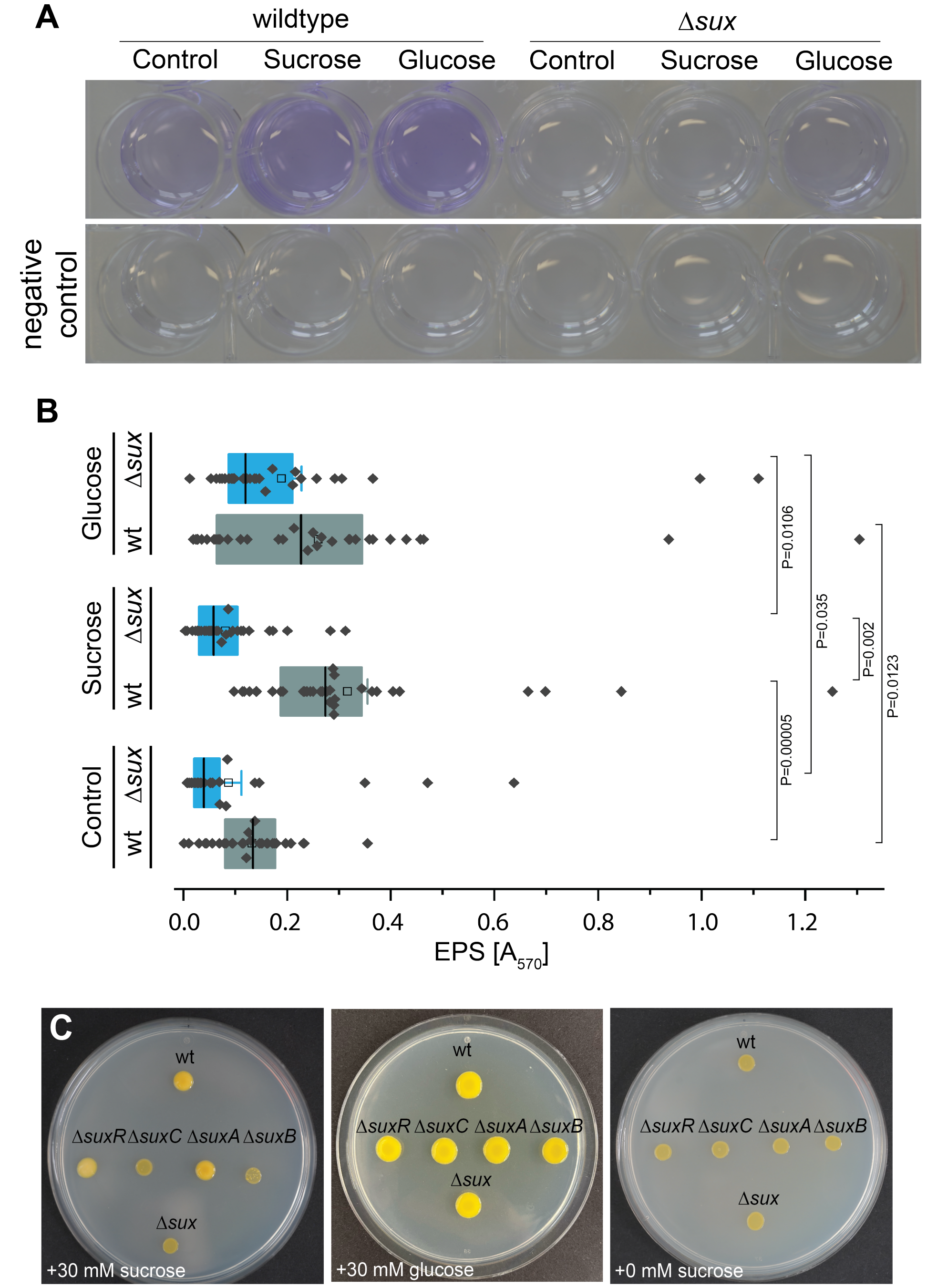


**Supplementary figure 5. Role of *sux* genes for EPS production and biofilm formation in glucose supplemented media. A)** Crystal violet staining of wildtype, Δ*sux* and negative control (no cells) in NB media ± 30 mM sucrose or glucose supply **B)** Quantification of crystal violet from wildtype and Δ*sux* in NB media ± 30 mM sucrose or glucose supply. Boxes range from 25th to 75th percentiles with median values shown as center lines and mean values as empty boxes with whiskers ± SEM. Significance between two groups was calculated using unpaired two-tailed Student`s t-test with 95 % confidence. P-values given for significant results. **C)** Representative colony phenotypes of wt and *sux* mutants in NB media supplemented with 30 mM sucrose, 30 mM glucose or without sugar supplementation.
